# Targeting default mode network connectivity with mindfulness-based fMRI neurofeedback: A pilot study among adolescents with affective disorder history

**DOI:** 10.1101/2022.08.22.504796

**Authors:** Jiahe Zhang, Jovicarole Raya, Francesca Morfini, Zoi Urban, David Pagliaccio, Anastasia Yendiki, Randy P. Auerbach, Clemens C.C. Bauer, Susan Whitfield-Gabrieli

## Abstract

Adolescents experience alarmingly high rates of major depressive disorder (MDD), however, gold-standard treatments are only effective for ~50% of youth. Accordingly, there is a critical need to develop novel interventions, particularly ones that target neural mechanisms believed to potentiate depressive symptoms. Directly addressing this gap, we developed a mindfulness-based fMRI neurofeedback (mbNF) for adolescents that targets default mode network (DMN) hyperconnectivity, which has been implicated in the onset and maintenance of MDD. In this proof-of-concept study, adolescents (*n* = 9) with a lifetime history of depression and/or anxiety were administered clinical interviews and self-report questionnaires, and then, each participant’s DMN and central executive network (CEN) were personalized using a resting state fMRI localizer. After the localizer scan, adolescents completed a brief mindfulness training followed by a mbNF session in the scanner wherein they were instructed to volitionally reduce DMN relative to CEN activation by practicing mindfulness meditation. Several promising findings emerged. First, mbNF successfully engaged the target brain state during neurofeedback; participants spent more time in the target state with DMN activation lower than CEN activation. Second, in each of the nine adolescents, mbNF led to significantly reduced within-DMN connectivity, which correlated with post-mbNF increases in state mindfulness. Last, a reduction of within-DMN connectivity mediated the association between better mbNF performance and increased state mindfulness. These findings demonstrate that personalized mbNF can effectively and non-invasively modulate the intrinsic networks known to be associated with the emergence and persistence of depressive symptoms during adolescence.

## INTRODUCTION

In the United States, major depressive disorder (MDD) results in over $200 billion of lost productivity and health care expenses each year (1), and for adolescents specifically, rates are alarmingly high (2). Gold-standard psychological and pharmacological treatments, however, are only effective for ~50% of youth (3), underscoring the critical need to develop novel treatments to improve clinical outcomes, particularly those that target core mechanisms that may underlie depression.

At the neural systems level, MDD is characterized by elevated resting state connectivity within the default mode network (DMN), which includes core midline hub regions in the subgenual anterior cingulate cortex (sgACC), medial prefrontal cortex (MPFC), and posterior cingulate cortex (PCC) (4,5). DMN hyperconnectivity, especially sgACC hyperconnectivity, is associated with symptom severity in depressed individuals (6,7) and characterizes children with elevated familial risk for depression (8). The DMN is thought to facilitate patterns of depressogenic, self-referential processing and a heightened focus on distressing emotional states (4,9–11); in MDD, it is theorized that dysregulation of the DMN by top-down control networks such as the central executive network (CEN), as evidenced by altered connectivity between the DMN and CEN (12,13), also contributes to heightened self-focus. As such, hyperconnectivity within the DMN has been linked to rumination (i.e., the tendency to perseverate about one’s symptoms), a common trait that contributes to depression onset, maintenance, and recurrence (14–16) as well as predicts cognitive therapy non-response and relapse (15,17).

As DMN connectivity is a promising biomarker of MDD (7,18), new interventions targeting DMN have recently been explored. For example, transcranial magnetic stimulation (TMS) targeting the dorsolateral prefrontal cortex (i.e., a CEN node that is anticorrelated with DMN) normalizes DMN connectivity and improves depressive symptoms in adults (19). TMS, however, relies on an external source of energy to stimulate brain tissue and alter brain connectivity. TMS can be invasive, as it is a targeted transcutaneous procedure (20). Accordingly, additional safety and efficacy evidence is warranted before routine clinical TMS can be used with children and adolescents. By contrast, mindfulness-based interventions are non-invasive and also can lead to decreased DMN activity (21–25) and connectivity (26) by leveraging an individual’s own cognitive resources to train attentional focus to the present moment. Mindfulness-based therapeutic approaches also reduce depression symptoms (27,28) and improve depression treatment outcomes (29,30). Our and others’ research have demonstrated that adolescents can apply mindfulness practices to reduce depression symptoms, including perceived stress (31,32). Yet, specific depressive symptoms (e.g., inattention, lack of energy, apathy, etc.) may prevent adolescents from more successfully integrating and applying mindfulness strategies in daily life.

To facilitate the acquisition and utilization of mindfulness strategies, we recently developed a novel mindfulness-based fMRI neurofeedback (mbNF; (33)) approach, which is a non-invasive technique that allows people to track and directly modulate brain function. To date, neurofeedback studies in depression have frequently involved mood-related tasks, such as negative emotion induction or valenced autobiographical memory recall (see review in (34)). By contrast, our mbNF targets DMN connectivity given known associations with mindfulness and MDD. In this 15-minute neurofeedback paradigm, people observe a schematic visual representation of their brain activity and practice mindfulness to volitionally reduce DMN activation relative to CEN activation. We previously used this paradigm in adults with schizophrenia and demonstrated that mbNF reduced DMN connectivity and led to symptom reduction postintervention (35). This mbNF method is non-invasive, optimizes implementation of mindfulness to target the DMN, and has enormous potential to translate skill acquisition outside of the scanner.

Building on our prior fMRI neurofeedback research (35,36), we tested mbNF in adolescents with a history of affective disorders as a proof-of-concept. First, we tested whether adolescents can successfully engage CEN more than DMN during mbNF. Second, we tested whether mbNF leads to reduced DMN connectivity and an associated increase in state mindfulness. Third, we tested whether reduced DMN connectivity accounted for the association between successful neurofeedback and increased state mindfulness.

## METHODS AND MATERIALS

### Participants and Procedure

Adolescents (*n* = 9; 18.8 ± 0.7 years; 17-19 years; 66.7% females) who previously completed scans for the Boston Adolescent Neuroimaging of Depression and Anxiety Human Connectome project (BANDA; (37,38)) were re-contacted, screened and enrolled in this proof-of-concept study. Sociodemographic and clinical characteristics are summarized in **Table 1**; all participants reported a lifetime history of MDD and/or anxiety disorders.

**Table 1.**
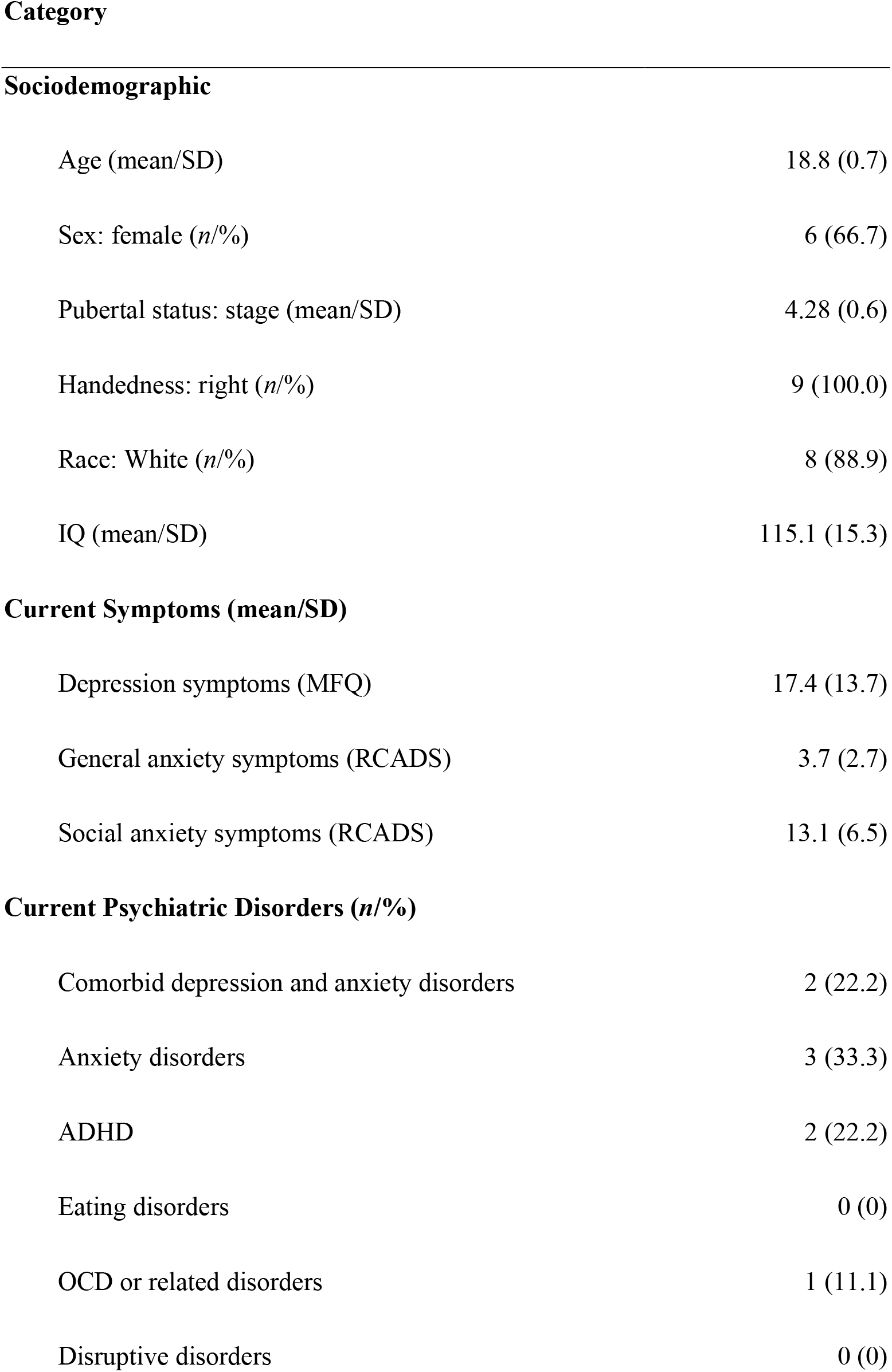
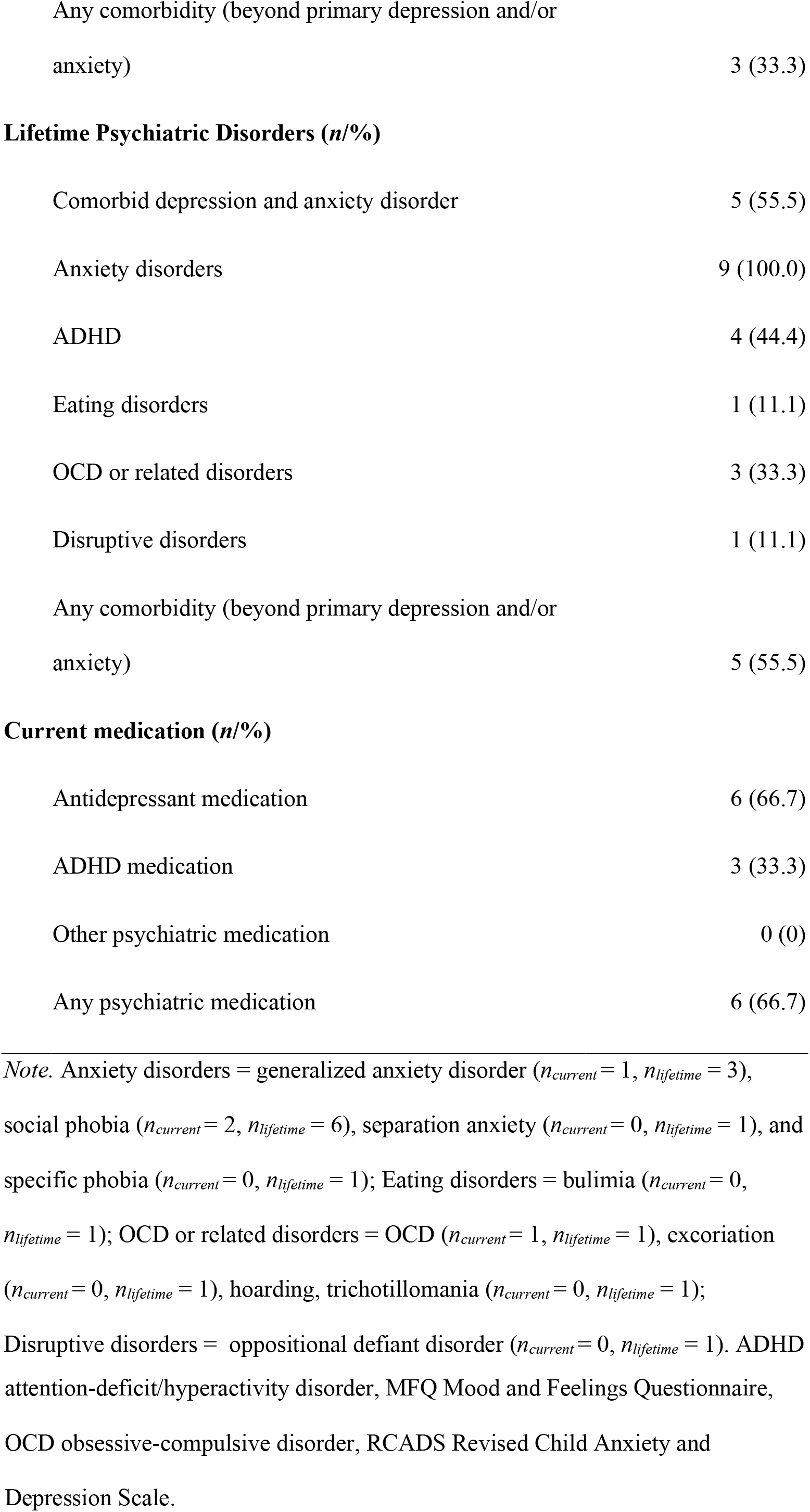
Sociodemographic and clinical information (*n* = 9) Category.

|  |  |
| --- | --- |
| Age (mean/SD) | 18.8 (0.7) |
| Sex: female ($n/\%$) | 6 (66.7) |
| Pubertal status: stage (mean/SD) | 4.28 (0.6) |
| Handedness: right ($n/\%$) | 9 (100.0) |
| Race: White ($n/\%$) | 8 (88.9) |
| IQ (mean/SD) | 115.1 (15.3) |
| **Current Symptoms (mean/SD)** |  |
| Depression symptoms (MFQ) | 17.4 (13.7) |
| General anxiety symptoms (RCADS) | 3.7 (2.7) |
| Social anxiety symptoms (RCADS) | 13.1 (6.5) |
| **Current Psychiatric Disorders ( $n/\%$ )** |  |
| Comorbid depression and anxiety disorders | 2 (22.2) |
| Anxiety disorders | 3 (33.3) |
| ADHD | 2 (22.2) |
| Eating disorders | 0 (0) |
| OCD or related disorders | 1 (11.1) |
| Disruptive disorders | 0 (0) |
| Any comorbidity (beyond primary depression and/or anxiety) | 3 (33.3) |
| **Lifetime Psychiatric Disorders (*n*/%)** |  |
| Comorbid depression and anxiety disorder | 5 (55.5) |
| Anxiety disorders | 9 (100.0) |
| ADHD | 4 (44.4) |
| Eating disorders | 1 (11.1) |
| OCD or related disorders | 3 (33.3) |
| Disruptive disorders | 1 (11.1) |
| Any comorbidity (beyond primary depression and/or anxiety) | 5 (55.5) |
| **Current medication (*n*/%)** |  |
| Antidepressant medication | 6 (66.7) |
| ADHD medication | 3 (33.3) |
| Other psychiatric medication | 0 (0) |
| Any psychiatric medication | 6 (66.7) |

*Note.* Anxiety disorders = generalized anxiety disorder ( $n_{current} = 1$ , $n_{lifetime} = 3$ ), social phobia ( $n_{current} = 2$ , $n_{lifetime} = 6$ ), separation anxiety ( $n_{current} = 0$ , $n_{lifetime} = 1$ ), and specific phobia ( $n_{current} = 0$ , $n_{lifetime} = 1$ ); Eating disorders = bulimia ( $n_{current} = 0$ , $n_{lifetime} = 1$ ); OCD or related disorders = OCD ( $n_{current} = 1$ , $n_{lifetime} = 1$ ), excoriation ( $n_{current} = 0$ , $n_{lifetime} = 1$ ), hoarding, trichotillomania ( $n_{current} = 0$ , $n_{lifetime} = 1$ ); Disruptive disorders = oppositional defiant disorder ( $n_{current} = 0$ , $n_{lifetime} = 1$ ). ADHD
attention-deficit/hyperactivity disorder, MFQ Mood and Feelings Questionnaire, OCD obsessive-compulsive disorder, RCADS Revised Child Anxiety and Depression Scale.

For Session 1, which was a follow-up to the BANDA protocol, study procedures were approved through the Mass General Brigham IRB. At the baseline visit, participants were administered a clinical interview and self-report assessments of depressive and anxiety symptoms. Then, each participant completed a localizer MRI session at the Athinoula A. Martinos Center for Biomedical Imaging. At the end of Session 1, participants were provided with information for Session 2 and interested participants were enrolled. Session 2 procedures were approved by the Northeastern University Institutional Review Board, and typically occurred within 2-3 weeks. Participants underwent mindfulness meditation training, completed a neurofeedback MRI session at the Northeastern University Biomedical Imaging Center, and completed pre- and post-scan state mindfulness assessments.

### Session 1

#### Clinical assessments

Participants were administered the Kiddie Schedule for Affective Disorders and Schizophrenia Present and Lifetime Version (KSADS; (39)) to assess the occurrence of psychiatric disorders since their prior study visit (i.e., past 2-3 years). Participants completed the Child Self-Report of the Mood and Feelings Questionnaire (MFQ; (40)) to assess depression symptom severity. The MFQ is a 33-item questionnaire, and the total score ranges between 0-66, with higher scores indicating more severe depression symptoms. Participants also completed the Revised Child Anxiety and Depression Scale (RCADS; (41)). The primary subscales of interest characterize general anxiety and social anxiety symptoms.

#### Functional Localizer

MRI data was acquired on a Siemens Prisma scanner with a 64-channel, phased-array head coil (Siemens Healthcare, Erlangen, Germany), including: a T1-weighted MPRAGE structural scan [0.8 mm isotropic voxel size, 208 slices, field-of-view (FOV) = 256 x 240 x 167 mm, repetition time (TR) = 2400 ms, echo time (TE) = 2.18 ms, flip angle (FA) = 8°] and two resting state fMRI scans (rs-fMRI: 5 minutes 46 seconds each, multiband acceleration factor = 8, 2 mm isotropic voxel size, 72 slices, FOV = 208 x 208 x 144 mm, TR = 800 ms, TE = 37 ms, FA = 52°) to identify participant-specific DMN and CEN maps.

Preprocessing of rs-fMRI data was performed in FSL 6.0 (42) and included: motion correction, brain extraction, co-registration, smoothing and bandpass filtering (see more details in (35)). We performed an independent components analysis (ICA) on the preprocessed functional scans using Melodic ICA version 3.14 (43) with dimensionality estimation using the Laplace approximation to the Bayesian evidence of the model. Each of the ~30 spatiotemporal components were statistically compared to atlas spatial maps of the DMN and CEN derived from rs-fMRI of ~1000 participants (44) using FSL’s “fslcc” tool and we select the ICA components that yielded the highest spatial correlation for each participant. These ICA components were thresholded to select the upper 10% of voxel loadings and then binarized to obtain participant-specific DMN and CEN masks to be used during neurofeedback in Session 2. Visual inspection was performed and all components maps were determined to be satisfactory in covering canonical DMN and CEN brain regions (45).

### Session 2

In Session 2, participants completed: pre-mbNF state mindfulness assessment, mindfulness training, structural scan, pre-mbNF rs-fMRI, mbNF, post-mbNF rs-fMRI, post-mbNF state mindfulness assessment.

#### State Mindfulness

Participants completed the State Mindfulness Scale (SMS; (46)) by indicating on a 5-point scale their perceived level of awareness and attention to their present experience during the last 15 minutes. The SMS was scored both as a sum of all 21 items (ranges 0-105) as well as two subscales assessing 15 items on mindfulness of the mind (i.e., thoughts and emotions; ranges 0-75) and 6 items on mindfulness of the body (i.e., movement and physical sensations; ranges 0-30).

#### Mindfulness Training

We trained participants on a mindfulness technique called “mental noting”. Mental noting is a major component of Vipassana or insight meditation practice (47) and consists of the factors “concentration” and “observing sensory experience”. The experimenter first explained to the participants that mental noting entails being aware of the sensory experience without engaging in or dwelling on the details of the content; in other words, one would “note” the sensory modality (e.g., “hearing”, “seeing”, “feeling”) at the forefront of their awareness and then let it go after it has been noted. The experimenter also introduced the concept of an “anchor”, or a sensory experience to which one could easily switch their attention, such as breathing. Participants were encouraged to use their personal “anchors” when they noted consecutive “thinking” (i.e., rumination). The experimenter demonstrated noting out loud by verbalizing the predominant sensory modality approximately once per second. Participants were then asked to practice mental noting out loud to demonstrate the ability to describe sensory awareness without engaging in the content and stop consecutive “thinking”.

To assess the effectiveness of mindfulness training, participants listened to audio recordings of brief stories before and after training and were asked to complete mental noting after training. This included five stories describing everyday, mundane characters and events recorded on audio in neutral tone. Each story lasted about half a minute and included 20 unique details. For example, in the sentence “Grandpa owns a garden”, there are 3 unique details (i.e., “grandpa”, “owns”, “garden”). Immediately before the participants learned about mental noting, they listened to one story and were asked to freely recall as many details as possible. After participants learned mental noting, we asked them to practice mental noting while we played stories (i.e., introducing salient auditory stimulus). The number of stories ranged between 2-4 and we stopped either after the participant was comfortable at the practice or after all remaining 4 stories were played. Compared to the baseline test when participants fully attended to story playback, usage of noting strategy during playback led to a significantly reduced number of details recalled [t(8) = −16.20, p < 0.001], indicating that participants were successfully engaging the noting strategy and therefore not retaining the details of the story.

#### mbNF

During the neurofeedback task, participants observed one centrally displayed white dot and two circles on the screen (**Fig.1**). The schema displays a red circle on top with a blue circle below. Participants were instructed to move the white dot into the red circle by performing mental noting. We explained that upward movement of the dot was associated with effective mental noting performance and downward movement was associated with excessive self-related processing and mind-wandering. We asked the participants immediately after the neurofeedback task whether they had used the mental noting strategy during neurofeedback; each participant reported using mental noting.

**Figure 1.**
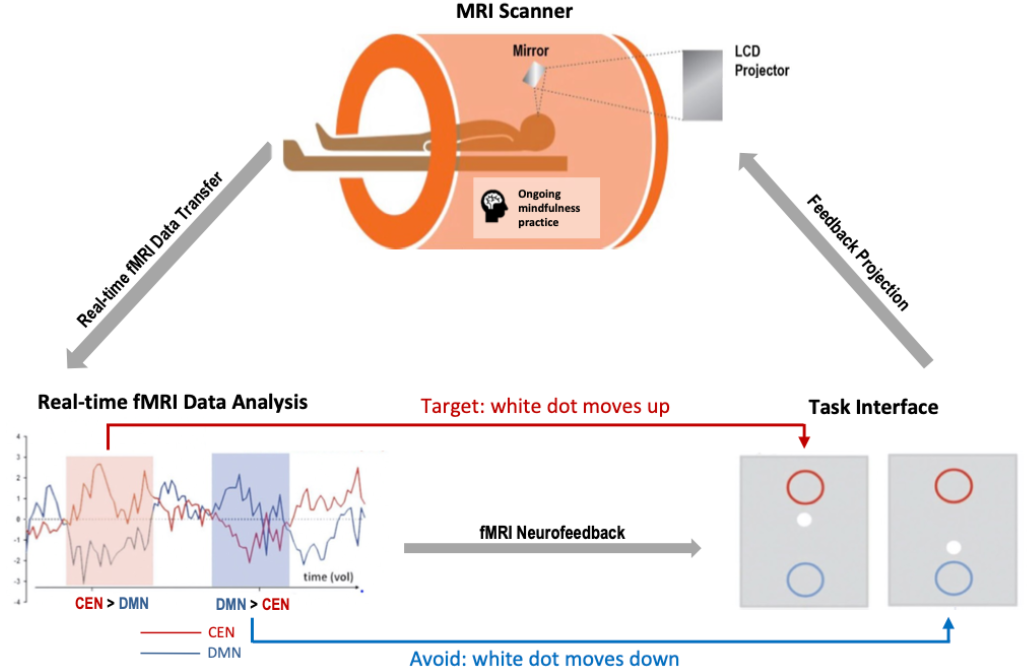
Mindfulness-based neurofeedback (mbNF). During mbNF, participants were instructed to practice mindfulness to move the white dot on the screen up into the red circle. The movement of the white dot was dependent on a real-time analysis of the fMRI data that computed the difference in personalized CEN and DMN activations. When CEN activation is higher than DMN activation, the white dot moves up; when CEN activation is lower than DMN activation, the white dot moves down.

There were five neurofeedback runs in this protocol and each lasted 2.5 minutes. We provided real-time feedback to the participants by means of a Positive Diametric Activity (PDA) metric (Bauer et al., 2019; Bauer et al., 2020) (**Fig. 1**). The PDA metric is based on the hypothesis that there is a causal neural mechanism by which the CEN negatively regulates the DMN (Chen et al., 2013). Accordingly, we defined the PDA as CEN activation estimate minus DMN activation estimate. The CEN and DMN estimates were calculated as the mean activity estimate across all voxels within each participant-specific mask as defined by Session 1 rs-fMRI (see *Session 1: Functional Localizer)* and co-registered to the current fMRI volumes. To accomplish the voxel-wise estimation in real-time (48), we first collected 30 seconds of baseline data and then continuously performed an incremental general linear model (GLM) fit with subsequent incoming images. This method accounts for the mean voxel signal and linear trends. To discount components of the voxel signal due to nuisance sources (e.g., low-frequency signal drifts), the GLM reconstruction of the expected voxel intensity at time *t* was subtracted from the measured voxel intensity at time *t*, leaving a residual signal that has components due to two sources: BOLD signal fluctuations and unmodeled fMRI noise. This residual was scaled by an estimate of voxel reliability, which was computed as the average GLM residual over the first 25 functional images of the baseline. This analysis resulted in an estimate of the strength of activation at each voxel at time *t* in units of standard deviation.

### Session 2 MRI Acquisition, Preprocessing and Data Analytic Overview

#### MRI Acquisition

A structural scan was acquired using a T1-weighted MPRAGE pulse sequence (1 mm isotropic voxel size, 176 slices, FOV = 256 x 256 x 256 mm, TR = 2530 ms, TE = 46 ms, FA = 7°). For functional images including during mbNF, the BOLD signal was measured using a T2* weighted gradient-echo, echo-planar imaging (EPI) pulse sequence (2 mm isotropic voxel size, 68 slices, FOV = 256 x 256 x 256 mm, TR = 1200 ms, TE = 30 ms, FA = 72°). Each neurofeedback run lasted 2 minutes and 30 seconds. Immediately before and after mbNF, two rs-fMRI scans (5 minutes each) were acquired.

#### MRI Preprocessing

Preprocessing was performed using *fMRIPrep* 21.0.0 (49,50), which is based on *Nipype* 1.6.1 (51,52). In short, common preprocessing steps were performed including realignment, co-registration, normalization, susceptibility distortion correction, segmentation of gray matter (GM), white matter (WM), cerebrospinal fluid (CSF) tissues, skull stripping, and confounds extraction. See Supplementary Information for a detailed description. Visual quality control was performed on each preprocessed run.

Preprocessed data and confound time series were imported into the CONN Toolbox v20.b (53) where outlier identification was performed with the Artifact Detection Tools (ART, www.nitrc.org/projects/artifact_detect). Volumes with global signal z > 5 and framewise displacement > 0.9mm compared to the previous frame were flagged as outliers. In addition, in-scanner mean motion was defined as the mean framewise displacement of the whole run (42) and calculated separately for pre- and post rtfMRI-NF runs. Rs-fMRI runs were spatially smoothed with a 6mm Gaussian kernel. A principal component analysis identified noise components from the WM and CSF following CONN’s implementation (54) of the aCompCorr method (55). During denoising, we regressed out the effect of the top 5 WM noise components, top 5 CSF noise components, 12 realignment parameters (3 translation, 3 rotation, and their first derivatives), linear drift and its first derivative effect, motion outliers, and applied a band-pass filter of 0.008 – 0.09 HZ.

#### Data Analytic Overview

Using the CONN toolbox (53), we performed functional connectivity analyses seeding the sgACC (8mm-radius sphere around MNI −2, 22, −16) (56). For baseline brain-behavior correlation analysis, we searched the whole brain for regions where connectivity with the seed correlated with MFQ scores at *p* < 0.001 (uncorrected). For functional connectivity change, we used SPM small volume correction to search midline DMN regions (MPFC and PCC nodes as defined in CONN toolbox DMN network) for voxels whose connectivity with the seed region changed significantly after mbNF and reported clusters that survived a FDR-corrected threshold of *q* = 0.05. Both analyses controlled for framewise displacement (57). We performed MPFC-seeded analyses as well (8mm-radius sphere around MNI −1, 53, −3) (58) and results can be found in the Supplementary Information.

## RESULTS

### DMN-Depressive Symptom Severity Association

In line with prior research (6,7), current depression symptom severity (MFQ) positively correlated (p < 0.001, uncorrected) with functional connectivity between the sgACC seed (**Fig. 2A**) and several regions, including the MPFC (**Fig. 2B**; 21 voxels; peak at MNI −6, 66, 18), right lateral temporal cortex (32 voxels; peak at MNI) and middle frontal gyrus (28 voxels; peak at MNI). Specifically, more severe depression symptoms were associated with greater connectivity between the sgACC seed and the MPFC (scatterplot shown in **Fig. 2C**).

**Figure 2.**
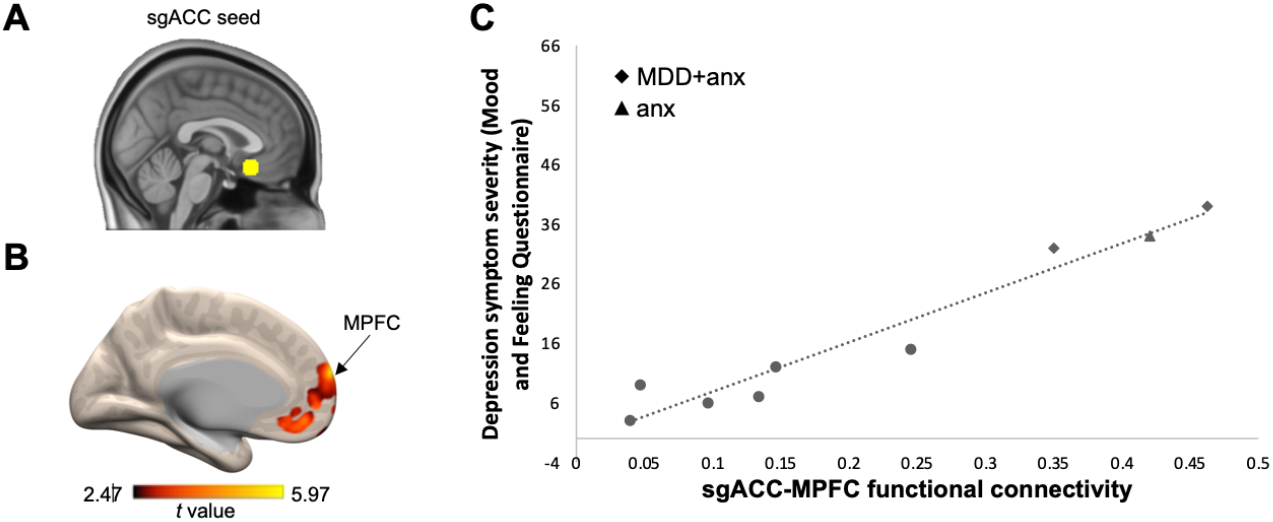
Higher DMN functional connectivity was associated with more severe depression symptoms at baseline. **A)** We used a 8mm spherical seed in the sgACC (56). **B)** Functional connectivity between the sgACC seed and the MPFC positively correlated with symptom severity. Higher MFQ score indicates higher severity. Arrow indicates the peak of the MPFC cluster that survived *p* < 0.001 (uncorrected). Figure is displayed at *p* < 0.05 (uncorrected) and the color bar range reflects minimum and maximum *t* values in the connectivity map. **C)** Scatterplot illustrates the correlation between baseline MFQ and baseline sgACC-MPFC functional connectivity. All participants had a lifetime history of MDD and/or anxiety. Patients with current diagnoses are labeled with a diamond for having comorbid anxiety and depression (‘MDD + anx’) and a triangle for having anxiety only (‘anx’).

### Neurofeedback Performance

Averaging across all 5 neurofeedback runs, participants spent more time in the target brain state (i.e., CEN > DMN activation) than expected by chance *(p* = .038, one-tailed t-test against 50% chance). Additionally, participants exhibited marginally higher CEN activation than DMN activation *(p* = .071, one-tailed paired-sample t-test), however, this effect was non-significant.

### Functional Connectivity Change Following mbNF

To test for changes in DMN functional connectivity following mbNF, we compared sgACC seed functional connectivity pre-vs. post-mbNF using a paired-sample t-test. We found that, at the group level, the sgACC seed showed significantly reduced functional connectivity (*q*_FDR_ < 0.05) to both MPFC (1003 voxels; MNI −8, 60, −6) and PCC (1185 voxels; MNI −4, −62, 28) after mbNF (**Fig. 3A**). Furthermore, when visualizing the individual-level data, we found that all nine participants showed sgACC-mPFC connectivity reduction (**Fig. 3B**). Additionally, we found a negative correlation between sgACC-MPFC connectivity change and neurofeedback performance. Participants who spent more time in the target brain state on the last neurofeedback run showed greater reduction in sgACC-MPFC functional connectivity (*r* = −.67, *p* = .048; **Fig. 3C**). Average time spent in target state across 5 neurofeedback runs did not correlate with functional connectivity change.

**Figure 3.**
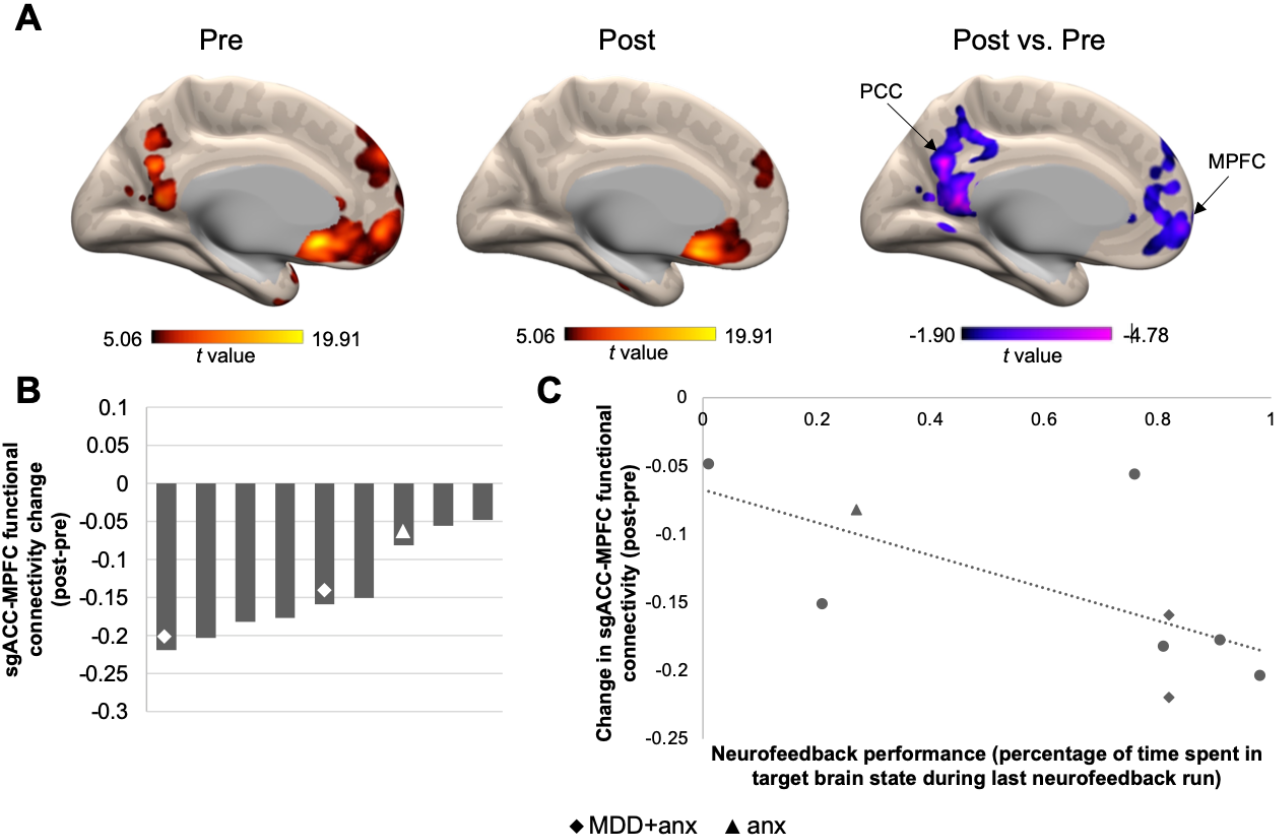
One session of mbNF reduced DMN functional connectivity. **A)** A t-test revealed that after mbNF, there was reduced connectivity between sgACC seed and midline DMN regions. Arrows indicate peaks in MPFC and PCC that survived *q*_FDR_ < 0.05. Pre and post connectivity maps are displayed at *p* < 0.001 (uncorrected). Post vs. Pre contrast map is displayed at *p* < 0.05 (uncorrected). Color bar ranges reflect minimum and maximum *t* values in the maps. **B)** Reduced sgACC-MPFC connectivity was found in all participants. Each bar represents the change in functional connectivity strength in a participant. **C)** Reduced sgACC-MPFC connectivity was associated with better neurofeedback performance. Patients with current diagnoses are labeled. All participants had a lifetime history of MDD and/or anxiety. Patients with current diagnoses are labeled with a diamond for having comorbid anxiety and depression (‘MDD + anx’) and a triangle for having anxiety only (‘anx’).

### Changes in State Mindfulness Pre- to Post-mbNF

Compared to pre-mbNF, participants reported significantly increased total state mindfulness after mbNF [*t*(8) = 1.90, *p* = .047], which was similarly observed in the mind [*t*(8) = 1.56, *p* = .079] and body subscales [*t*(8) = 2.26, *p* = .027]. As hypothesized, change in state mindfulness was positively correlated with neurofeedback performance. Specifically, participants who spent more time in the target brain state on the final neurofeedback run showed greater increases in SMS total (*r* = .69, *p* = .039) as well as both the mind [*r* = .71, *p* = .031] and body subscales [*r* = .59, *p* = .093]. Further, we found a negative correlation between change in functional connectivity and change in state mindfulness. Relative to pre-mbNF, more reduction in sgACC-MPFC functional connectivity was associated with greater increases in state mindfulness (SMS Total) following mbNF (**Fig. 4A**; *r* = −.88, *p* = .002). This association was consistent across the subscales (**Supplementary Fig. 1**; SMS Mind: *r* = −.87, *p* =.002; SMS Body: *r* = −.82, *p* = .007).

**Figure 4.**
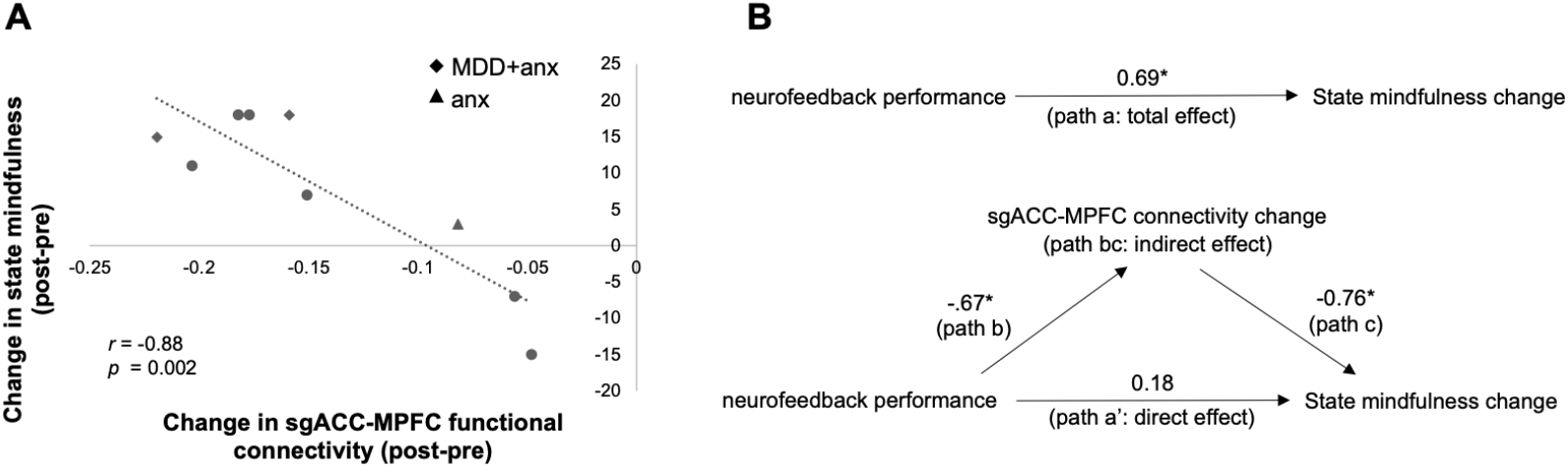
One session of mbNF induced state mindfulness change. **A)** Higher increase in state mindfulness after mbNF was associated with more decrease in sgACC-MPFC functional connectivity. Patients with current diagnoses are labeled. Anx: anxiety; dep: depression. **B)** Reduction in sgACC-MPFC connectivity fully mediated the association between better neurofeedback performance and increase in state mindfulness. Arrows indicate paths and path values indicate standardized beta weights. The upper panel shows the total effect (unmediated path a, total effect) from neurofeedback performance to state mindfulness change. In the lower panel, the effect of neurofeedback performance on state mindfulness change is fully mediated by the change in sgACC-MPFC functional connectivity. The direct effect of neurofeedback performance to state mindfulness change is indicated by path a’ and the indirect effect is indicated by the bc path (i.e., path b*path c). *\*p* < 0.05.

### DMN Functional Connectivity as a Mediator

Using mediation analysis (59,60), we found that functional connectivity change partially mediated the association between neurofeedback performance and state mindfulness change. We first regressed state mindfulness change on neurofeedback performance [*b* = 23.22, *β* = 0.69, *t* = 2.53, *p* = .039] (**Fig. 4B**; total effect, path a) and sgACC-MPFC connectivity change on neurofeedback performance *[b* = −0.12, *β* = −0.67, *t* = −2.39, *p* = .048] (**Fig. 4B**; path b). Controlling for neurofeedback performance, the mediator (sgACC-MPFC connectivity change) significantly predicted state mindfulness change [*b* = −141.75, *β* = −0.76, *t* = −3.07, *p* = .022] (Fig. 4B; path c). Further, controlling for the mediator (sgACC-MPFC connectivity change), neurofeedback performance was no longer a significant predictor of state mindfulness change [*b* = 6.07, *β* = 0.18, *t* = 0.73, *p* = .493] (**Fig. 4B**; direct effect, path a’). The Sobel test indicated that mediation by the indirect effect (path bc) was approaching significance (*t* = 1.88, *p* = .060).

## DISCUSSION

Depression is one of the most common mental disorders among adolescents, resulting in severe impairments. Accordingly, there is an urgent need to develop novel treatments to address escalating rates of depression among adolescents. In this proof-of-concept study, several key findings emerged, which support the feasibility of using fMRI neurofeedback to target adolescent depression symptoms. First, we demonstrated that participants successfully activated their individualized CEN more than DMN during mbNF. Second, we found that one session of successful mbNF led to reduced DMN connectivity and increased state mindfulness. Last, we found that reduction in sgACC-MPFC connectivity mediated the association between better neurofeedback performance and increase in state mindfulness.

The DMN consists of an ensemble of regions whose hyperactivation and hyperconnectivity result in altered self-referential processing and for some, may lead to rumination commonly occurring in MDD (7,18). There are a number of neuroanatomical landmarks implicated in the DMN (e.g., MPFC, PCC, angular gyrus, etc.) and our seed-based analysis approach privileged the sgACC as it shows specific alterations in MDD, including reduced glial cell count (61), abnormal blood flow and metabolism (62,63), abnormal thickness (64,65), reduced volume (61,62,66), and structural connectivity (67–69). Consistent with previous research, our sgACC-based functional connectivity findings suggest that the sgACC is a special hub in depression pathology and treatment. Similar to previous studies (6–8), we found that at baseline (i.e., prior to mbNF), elevated sgACC connectivity to other DMN nodes was associated with more severe depression symptoms.

Our study also provided more causal evidence that after one session of mbNF, reduced DMN connectivity partially mediated the association between better neurofeedback performance and increased state mindfulness. This suggests that a change in DMN connectivity was necessary to facilitate the behavioral change (i.e., state mindfulness) post-mbNF and is consistent with previous literature showing association between higher trait mindfulness and lower DMN connectivity (70,71). In other words, dampening of DMN activity during mbNF and reduced DMN connectivity post-mbNF may provide favorable conditions for mindfulness acquisition. In a similar vein, a recent study teaching healthy adolescents to regulate PCC activity using mindfulness meditation demonstrated increased mindfulness immediately after the training as well as after one week (72). In both studies, participants down-regulated DMN activity with help from real-time neurofeedback and subsequently showed improvement in mindfulness, which may be a pathway to reduce negative repetitive thinking and other depression symptoms. These findings indicate that DMN-targeted real-time neurofeedback is a feasible approach to boost mindfulness acquisition and may help youth seeking to reap the benefits associated with mindfulness-based interventions overcome challenges that impede learning, such as training length (73) and existing or emerging mental health symptoms (74).

One major innovation in our study is that instead of targeting a single brain region (i.e., the PCC), we implemented network-based modulation (i.e., the DMN and the CEN). As cognitive neuroscience has evolved from localized models to attributing function to distributed systems and their interactions, neurofeedback research has expanded from feedback targeting activation of a single region to include feedback focused on multiple, related regions (75). In clinical settings, this may lead to more effective intervention because depression, like many other psychiatric disorders with heterogeneous phenotypes, shows networked neuropathology (76). By modulating entire circuitries, we may have a better chance of directly normalizing altered brain connections and their associated functions. This is also one reason that we believe circuitry-based real-time fMRI neurofeedback may outperform focally targeted methods such as TMS and ultrasound. Indeed, another recent real-time fMRI study targeting dorsolateral prefrontal cortex-precuneus circuitry led to reduced depressive symptoms in patients (77).

One of the most encouraging findings from our study is that all 9 individuals who participated in the mbNF protocol demonstrated reduced DMN connectivity. This may be attributed to the personalized design of the protocol where the neurofeedback targets (i.e., DMN and CEN) were individually and functionally localized for each participant, which is in line with the broader mission in psychiatry towards utilizing biomarkers to guide precision medicine (78)._By comparison, randomized controlled clinical trials for TMS – where treatment target is typically not functionally localized – demonstrates a response rate between 15%-37% and a remission rate between 14%-30% (79). Our proof-of-concept study is not a clinical trial, and nor did we test clinical outcomes. However, it may be that personalizing treatment, which is common in deep brain stimulation (80), may foster improved clinical outcomes for MDD.

The current pilot study should be considered an early-phase exploratory trial to test the feasibility of the mbNF approach and thus a single-group design was justified (81). Future studies should include appropriate control conditions as well as address other limitations in the current study, such as small sample size and lack of post-mbNF assessment of depression symptoms. Longitudinal symptom followup is critical for determining the trajectory of clinical benefit, as suggested by studies that indicate that the greatest symptom improvement may happen weeks to months after neurofeedback intervention (82). We can also further personalize the protocol by fine-tuning the optimal session length and number of sessions for each subject. For example, this study revealed that neurofeedback performance on the last run, not the average, was more predictive of neural and behavioral change, suggesting the length of the mbNF session may vary depending on how fast a participant reaches a certain threshold of performance. The scalability of the mbNF paradigm will also be greatly improved if this protocol can be implemented in less costly systems such as electroencephalography or functional near-infrared spectroscopy. In the long term, we also aim to develop a closed-loop system for delivering mbNF intervention when suboptimal brain states (e.g., ruminative or suicidal) are detected in patients.

In summary, the mbNF protocol is a non-invasive and personalizable tool that could offer early intervention and alleviate depression in adolescents. Building on these promising findings, a key next step is to determine whether this approach leads to improvements in depressive symptoms, which has enormous potential to revolutionize our approach to clinical care.

## Acknowledgements

This study was supported by the NIMH R56MH121426 (SWG, AY, RPA) and R61/R33MH113751-01A1 (SWG), the Poitras Center for Psychiatric Disorders Research at Massachusetts Institute of Technology (SWG), and the Tommy Fuss Fund (RPA). The content is solely the responsibility of the authors and does not necessarily represent the official views of the National Institutes of Health. We would like to acknowledge Dr. Zhenghan Qi and Anqi Hu for providing short stories that we adapted for mindfulness training assessment.

