## Supplementary Information for "Targeting default mode network connectivity with mindfulness-based fMRI neurofeedback: A pilot study among adolescents with affective disorder history"

### 1) Supplementary Methods: MRI Preprocessing

*Preprocessing of $B_0$ inhomogeneity mappings*. A $B_0$-nonuniformity map (or *fieldmap*) was estimated based on two (or more) echo-planar imaging (EPI) references with topup (1).

*Anatomical data preprocessing*. The T1-weighted (T1w) image was corrected for intensity non-uniformity (INU) with N4BiasFieldCorrection (2), distributed with ANTs 2.3.3 (3), and used as T1w-reference throughout the workflow. The T1w-reference was then skull-stripped with a *Nipype* implementation of the antsBrainExtraction.sh workflow (from ANTs), using OASIS30ANTs as target template. Brain tissue segmentation of cerebrospinal fluid (CSF), white-matter (WM) and gray-matter (GM) was performed on the brain-extracted T1w using fast (FSL 6.0.5.1:57b01774, RRID:SCR_002823) (4). Brain surfaces were reconstructed using recon-all (FreeSurfer 6.0.1, RRID:SCR_001847) (5), and the brain mask estimated previously was refined with a custom variation of the method to reconcile ANTs-derived and FreeSurfer-derived segmentations of the cortical gray-matter of Mindboggle (RRID:SCR_002438) (6). Volume-based spatial normalization to a standard space (MNI152NLin6Asym) was performed through nonlinear registration with antsRegistration (ANTs 2.3.3), using brain-extracted versions of both T1w reference and the T1w template. The following templates were selected for spatial normalization: *FSL's MNI ICBM 152 non-linear 6th Generation Asymmetric Average Brain Stereotaxic Registration Model* (RRID:SCR_002823; TemplateFlow ID: MNI152NLin6Asym) (7).

*Functional data preprocessing.* For each of the 4 BOLD resting state runs, the following preprocessing was performed. First, a reference volume and its skull-stripped version were generated using a custom methodology of *fMRIPrep*. Head-motion parameters with respect to the BOLD reference (transformation matrices, and six corresponding rotation and translation parameters) are estimated before any spatiotemporal filtering using mcflirt (FSL 6.0.5.1:57b01774) (8). The estimated *fieldmap* was then aligned with rigid-registration to the target EPI (echo-planar imaging) reference run. The field coefficients were mapped on to the reference EPI using the transform. The BOLD reference was then co-registered to the T1w reference using bbregister (FreeSurfer) which implements boundary-based registration (9). Co-registration was configured with six degrees of freedom. Several confounding time-series were calculated based on the *preprocessed BOLD*, including framewise displacement (FD) and DVARS. FD was computed using two formulations following Power (absolute sum of relative motions) (10) and Jenkinson (relative root mean square displacement between affines) (11). FD and DVARS are calculated for each functional run, both using their implementations in *Nipype*, following the definitions by (10). The BOLD time-series were resampled into standard space, generating a *preprocessed BOLD run in MNI152NLin6Asym space*. First, a reference volume and its skull-stripped version were generated using a custom methodology of *fMRIPrep*. All resamplings can be performed with *a single interpolation step* by composing all the pertinent transformations (i.e. head-motion transform matrices, susceptibility distortion correction when available, and co-registrations to anatomical and output spaces). Gridded (volumetric) resamplings were performed using antsApplyTransforms (ANTs), configured with Lanczos interpolation to minimize the smoothing effects of other kernels (12). Non-gridded (surface) resamplings were performed using mri_vol2surf (FreeSurfer). Many internal operations of *fMRIPrep* use *Nilearn* 0.8.1 (RRID:SCR_001362) (13), mostly within the functional processing workflow.

**2) Supplementary Results: MPFC-Seeded Connectivity.**

A whole-brain analysis seeded in the MPFC (14) did not yield significant clusters that correlated with MFQ scores at baseline. We found significantly reduced functional connectivity ($p < 0.001$, uncorrected) to multiple DMN nodes post-mbNF, including the PCC, angular gyrus and lateral temporal cortex. MPFC-PCC functional connectivity change, however, did not correlate with state mindfulness change.

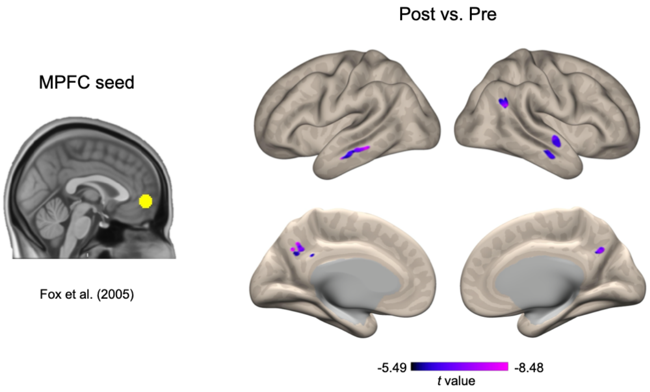


**3) Supplementary Figure 1.** One session of mbNF induced state mindfulness changes on both body and mind subscales. Higher increase in both body and mind mindfulness after mbNF was associated with more decrease in sgACC-MPFC functional connectivity. Patients with current diagnoses are labeled. Anx: anxiety; dep: depression.

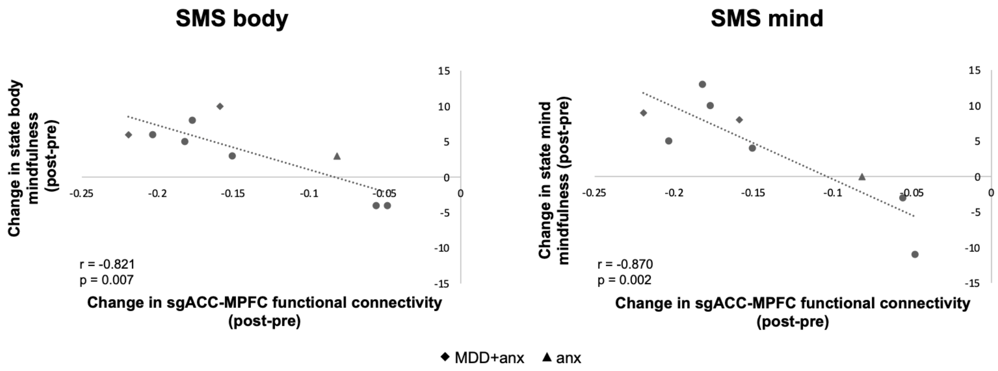
